# *Bifidobacterium longum* subsp*. longum* BG-L47 boosts growth and activity of *Limosilactobacillus reuteri* DSM 17938 and its extracellular membrane vesicles

**DOI:** 10.1101/2024.02.12.579962

**Authors:** Ludwig Ermann Lundberg, Punya Pallabi Mishra, Peidi Liu, Manuel Mata Forsberg, Eva Sverremark-Ekström, Gianfranco Grompone, Sebastian Håkansson, Caroline Linninge, Stefan Roos

**Affiliations:** Department of Molecular Sciences, Uppsala BioCenter, Swedish University of Agricultural Sciences, Uppsala, Sweden; BioGaia, SE-103 64 Stockholm, Sweden; MetaboGen, SE-411 26, Gothenburg, Sweden; The Department of Molecular Biosciences, The Wenner-Gren Institute, Stockholm University, Stockholm, Sweden; Division of Applied Microbiology, Department of Chemistry, Lund University, Lund, Sweden; Department of Food Technology, Engineering and Nutrition, Lund University, Lund, Sweden

**Author notes:** **Address for correspondence:** Ludwig Ermann Lundberg, Department of Molecular Sciences, Uppsala BioCenter, Swedish University of Agricultural Sciences, Uppsala, Sweden.

## Abstract

The aim was to identify a *Bifidobacterium* strain that improved the performance of *Limosilactobacillus reuteri* DSM 17938. Initial tests showed that *Bifidobacterium longum* subsp. *longum* strains boosted the growth of DSM 17938 during *in vivo-*like conditions. Further characterization revealed that one of the strains, BG-L47, had better bile and acid tolerance compared to BG-L48, as well as mucus adhesion compared to both BG-L48 and the control strain BB536. BG-L47 also had the capacity to metabolize a broad range of carbohydrates and sugar alcohols. Mapping of glycoside hydrolase (GH) genes of BG-L47 and BB536 revealed many GHs associated with plant-fiber utilization. However, BG-L47 had a broader phenotypic fiber utilization capacity. In addition, *B. longum* subsp. *longum* cells boosted the bioactivity of extracellular membrane vesicles (MV) produced by *L. reuteri* DSM 17938 during cocultivation. Secreted 5’ nucleotidase (5’NT), an enzyme that converts AMP into the signal molecule adenosine, was increased in MV boosted by BG-L47. The MV exerted an improved antagonistic effect on the pain receptor TRPV1 and increased the expression of the immune development markers IL-6 and IL-1ß in a PBMC model. Finally, the safety of BG-L47 was evaluated both by genome safety assessment and in a human safety study. Microbiota analysis showed that the treatment did not induce significant changes in the composition. In conclusion, *B. longum* subsp. *longum* BG-L47 has favorable physiological properties, can boost the *in vitro* activity of *L. reuteri* DSM 17938, and is safe for consumption, making it a candidate for further evaluation in probiotic studies.

**Importance:** By using probiotics that contain a combination of strains with synergistic properties, the likelihood of achieving beneficial interactions with the host can increase. In this study, we first performed a broad screening of *Bifidobacterium longum* subsp. *longum* strains in terms of synergistic potential and physiological properties. We identified a superior strain, BG-L47, with favorable characteristics and potential to boost the activity of the known probiotic strain *Limosilactobacillus reuteri* DSM 17938. Further, we demonstrated that BG-L47 is safe for consumption in a human randomized clinical study and by performing a genome safety assessment. This work illustrates that bacteria-bacteria interactions differ at the strain level and further provides a strategy for finding and selecting companion strains of probiotics.

## Introduction

In nature, microorganisms usually coexist in complex ecosystems (1), allowing for many potential interactions that can be either synergistic, neutral, or antagonistic (2). Similarly, increased efficacy of a probiotic bacterium can potentially be achieved by co-producing or co-administering it with partner strains. Synergistic and mutualistic interactions between the strains can potentially give rise to increased tolerance and ecological fitness, as well as potentiate host development and shape host fitness, ultimately improving interactions with the host (3–5). Olsson and colleagues demonstrated that *B. longum* stabilizes the human microbiota, which underlines that bacterial interactions in the intestine can have implications for human health (6).

Synergistic effects among lactobacilli and bifidobacteria have previously been investigated. Zhuge and colleagues (7) demonstrated that *Lactobacillus salivarius* and *Bifidobacterium longum* subsp. *longum* (*B. longum* subsp. *longum* is hereafter referred to as *B. longum* unless another subspecies is specified) acted synergistically to reduce proinflammatory cytokines and alleviate gut dysbiosis in rats with experimentally induced liver injury. Combination treatment using both *Lactobacillus acidophilus* and *Bifidobacterium animalis* subsp. *lactis* has been successful in relieving symptoms of bloating in individuals with functional bowel disorders (8). Cheng and colleagues showed that *Bifidobacterium breve* can provide *Limosilactobacillus reuteri* with 1,2-propanediol that *L. reuteri* can use as an electron acceptor, ultimately resulting in better energy yield during growth (9). Other approaches, such as evaluation of the combined protective effect of conditioned medium from *Lactobacillus acidophilus*, *B. longum* subsp. *infantis*, and *Lactobacillus plantarum* on necrotizing enterocolitis in a rodent model have also been tested. It was demonstrated that the incidence of intestinal injury was significantly reduced in a synergistic manner by the added supernatants (10). Synergy between separately cultivated lactobacilli and bifidobacteria was demonstrated by Li et al. (2019) who observed a greater anti-inflammatory effect in HT-29 cells when they combined the two bacteria (11). Furthermore, in a randomized controlled trial (RCT) in patients with cystic fibrosis, a fecal microbiota dominated by *Bifidobacterium* was more likely to occur in patients receiving a strain of *Lacticaseibacillus rhamnosus* (12).

*L. reuteri* DSM 17938 is a well-studied probiotic strain that has shown positive clinical effects in alleviating various disorders such as constipation, infantile colic, and functional abdominal pain (13–17). The mechanisms by which *L. reuteri* mediates these effects are largely unknown but are likely to be multifactorial in nature. However, several preclinical observations have been linked to the probiotic action of *L. reuteri* DSM 17938: e.g. exerting a modulatory effect on the immune system (18), enhancing the epithelial integrity (19), antagonizing pain receptor signaling via the pain receptor Transient receptor potential vanilloid 1 (TRPV1) (20) and affecting gut motility (21).

Bifidobacteria are early colonizers of the human gut and are often used in probiotic products (22). Continuously accumulating evidence highlights bifidobacteria as important inhabitants of the gut throughout life (23,24). They are known stimulators of the immune system and producers of short-chain fatty acids (SCFA) and other metabolites, which together contribute to intestinal and immune homeostasis (25). Bifidobacteria can also stimulate the growth of other microorganisms (23,26). Early colonization of *Bifidobacterium* strains is important for the infant immunological maturation (27). Furthermore, disturbance or absence of *Bifidobacterium* in the gut has been associated with elevated levels of intestinal inflammation markers (28), as well as with diseases (29) such as coeliac disease (30) and allergic dermatitis (31). So naturally occurring bifidobacteria are undeniably important members of the microbiota, but this group of bacteria has also shown promising probiotic capability. For example, supplementation with *B. longum* subsp*. infantis* EVC001 showed potential to correct systemic inflammation and altered immune regulation by silencing intestinal T helper 2 and T helper 17 as well as inducing IFN-ß in infants (28). Fang et. al have demonstrated that *B. longum* CCFM1029 can upregulate tryptophan metabolism and produce indole-3-carbaldehyde, which activated the aryl hydrocarbon receptor(ahr)-mediated immune response and alleviated atopic dermatitis symptoms (32). Products with bifidobacteria have also been successfully used as agents to treat diarrhea (33) and infantile colic (34).

In recent decades, it has been revealed that bacteria release extracellular membrane vesicles (MV) which are potent mediators of various types of signals (35). While MV from pathogens may carry virulence factors or toxins, there is increasing evidence that MV derived from mutualistic and probiotic bacteria exhibit beneficial effects on the host and participate in the probiotic action (36); (37,38). *L. reuteri* DSM 17938 also produces MV (39) that can modulate intestinal motility in an *ex vivo* mice model (40) and exhibit immunomodulatory effects in a PBMC model (41). Furthermore, we recently demonstrated the multifunctionality of these vesicles, including attenuation of immune responses in PBMCs, antagonism of TRPV1 in dorsal root ganglia, and protection of epithelial cells from ETEC-induced leakage (42). It was also shown that the MV from *L. reuteri* DSM 17938 and BG-R46 carry a surface-anchored 5’-nucleotidase (5’NT) that can convert AMP to the signaling molecule adenosine. Accordingly, Liu et al. 2019 (43) showed that new-born mice supplemented with *L. reuteri* had increased levels of adenosine in plasma after treatment. Additionally, Liu et al 2023 showed that DSM 17938 and BG-R46 can prolong the survival of scurfy mice by anti-inflammatory adenosine signaling (44).

Thus, strains of both *L. reuteri* and *B. longum* can be potent regulators of gastrointestinal functions and have been described to mediate multiple health-promoting effects. We sought to find a strain of bifidobacteria that enhances the probiotic activity of *L. reuteri* DSM 17938. To reduce the number of strains to evaluate, we initially screened several strains of *Bifidobacterium* for their basic properties and selected two *B. longum* strains for further characterization in search of a synergistic partner strain. The strains were benchmarked against the well-studied *B. longum* strain BB536 by assessing the growth-stimulating effect on *L. reuteri* DSM 17938 and evaluating their basic probiotic characteristics and carbohydrate utilization capacity. One strain, BG-L47, was selected and its ability to stimulate the bioactivity of DSM 17938-derived MV was evaluated in several models. BG-L47 was finally evaluated from a safety perspective, both by assessment of the genome sequence, and in a clinical safety and tolerability study.

## Methods

### Strains and strain development

Multiple *Bifidobacterium* isolates were previously isolated from infant (15 days old) feces (45) of which two were selected for this study (Supplementary table 1). These bifidobacteria were modified through a selection procedure in which they were repeatably subcultured from single colonies until culture reproducibility and homogeneity in appearance and behavior were obtained. The strains were identified by 16S rRNA gene sequencing and subspecies-specific PCR (see below) and named *B. longum* subsp. *longum* BG-L47™ and BG-L48™ (trademarks of BioGaia AB, GenBank accession numbers MZ411576 and MZ411575). After being deposited to the German Collection of Microorganisms and Cell Cultures (DSMZ), the strains were designated DSM 32947 and DSM 32948. The mother strain of BG-L47, *B. longum* subsp. *longum* CF15-4C, was used for genetic comparisons. The above-mentioned strains were obtained from BioGaia AB (Stockholm, Sweden). *B. longum* subsp. *longum* strain BB536 (https://morinagamilk-ingredients.com/probiotics/bb536/) was isolated from infant feces in 1969 and has been used in probiotic products since 1971. BB536 was used as a comparison strain because it is a well-studied probiotic strain of *B. longum*. The strain was obtained from Arla AB (Stockholm, Sweden).

### PCR for identification of *Bifidobacterium longum* subspecies

Subspecies-specific PCR for differentiation between *B. longum subsp. longum* and *B. longum subsp. infantis* was performed. Primer pairs used were inf_2348_F (ATA CAG AAC CTT GGC CT) and inf_2348_R (GCG ATC ACA TGG ACG AGA AC) for subsp. *infantis* detection, and lon_0274_F (GAG GCG ATG GTC TGG AAG TT) and lon_0274_R (CCA CAT CGC CGA GAA GAT TC) for subsp. *longum* detection (46). The PCR program was 95°C, 5 min; 35x (95°C, 30 s; 62°C, 30 s; 72°C, 30 s), 72° C, 10 min, and held at 16°C. *B. longum* subsp. *longum* BB536 and *B. longum* subsp. *infantis* CCUG 30512B were used as controls. The PCR products were visualized by standard agarose gel electrophoresis.

### Growth stimulation of *L. reuteri* by co-cultivation with *B. longum*

*L. reuteri* DSM 17938 was coincubated with the bifidobacterial strains. The *B. longum* strains were grown anaerobically at 37° C in 10 ml MRS (Oxoid, Basingstoke, Hampshire, United Kingdom) for 42-48 h and *L. reuteri* DSM 17938 was cultivated in 10 ml MRS at 37° C for 18 h. The bacteria were thereafter centrifuged at 10 000 x *g* for 10 min, followed by resuspension in an equal volume of PBS. In the following cultivation, 2 ml of *L. reuteri* DSM 17938 suspension was inoculated into 400 ml of substrate and the volume of bifidobacterial suspension was 16 ml (4%), 40 ml (10%) or 100 ml (25%). The cultures were incubated anaerobically for 48 h. For this co-cultivation of the two types of bacteria, we used a simple substrate that simulate the conditions in the upper part of the intestine, Simulated Intestinal Medium (SIM), containing glucose as carbon source and that has a relatively low nutrient content and contains bile (Supplementary Table 2). *L. reuteri* DSM 17938 grown in SIM supplemented with 15 mM fructose was used as a control in all experiments. The co-cultured bacteria were analyzed on Homofermentative-Heterofermentative Differential (HHD) diagnostic agar plates(47), Bifidus selective medium (BSM) agar (Sigma) and MRS + mupirocin (50 μg/ml) to quantify the presence of bifidobacteria. Plates were incubated anaerobically for 48 h at 37° C.

### High Performance Liquid Chromatography

Acetate production was analyzed by high performance liquid chromatography (HPLC) using an Agilent 1100 Series (Agilent, Santa Clara, CA, USA), with a refractive index detector and an ion exclusion column (Rezex ROA - Organic Acid H+, 300 x 7.80 mm, Phenomenex). The mobile phase consisted of 5 mM H_2_SO_4_; flow rate of 0.6 ml min^-1^. Samples were prepared by mixing 700 µl of sample with 70 µl of 5 M H_2_SO_4_ before being centrifuged at 14 000 x *g* for 15 minutes. The supernatants were filtered through a 0.2 µm syringe filter into HPLC glass vials. Two internal references containing approximately 0.5 g/L and 4 g/L of acetate were used.

### NMR spectroscopy

The BG-L47 supplemented SIM medium was evaluated by nuclear magnetic resonance (NMR) where 600 µl was transferred to 5 mm NMR tubes and spectra were recorded on a Bruker Avance III 600 MHz spectrometer (Bruker Biospin, Billerica, MA, USA) with a 5 mm broadband observe detection Smartprobe equipped with z-gradient. The obtained 1H spectra were compared to the spectra of known electron acceptors of *L. reuteri* DSM 17938 including fructose, glycerol, pyruvate, citrate, nitrate, oxygen and 1,2-propanediol using a local database with the software ChenomX and data collected from the online database Human Metabolome Database (https://hmdb.ca). TopSpin, version 4.0.9 (Bruker) was used for data processing.

### Characterization of *B. longum* strains

#### Scanning electron microscopy

The scanning electron microscopy was performed at the centre for cellular imaging, Core Facilities, the Sahlgrenska Academy, University of Gothenburg in Sweden. Bacteria were lyophilized as described in Pang et al 2022 (42). Lyophilized bacteria were rehydrated in 0.1 M PBS for 20 min. Coverslips were coated with 0.1% poly-L-lysine for 10 minutes followed by drying for 5 min, after which they were incubated in cell suspension for 30 min at room temperature. Excess medium was removed, and the cells were washed and allowed to incubate in 2.5% glutaraldehyde in 0.1 M PIPES for 30 min at room temperature. The samples were post-fixated in 1% osmium tetroxide in 0.1 M PIPES for one hour at 4°C in the dark. After post-fixation, the samples were successively dehydrated in a series of ethanol solutions with increasing ethanol concentration (35-100%). The samples were suspended in HMDS solution and then allowed to air dry. The samples were sputtered with gold (Emitech, Taunusstein, Germany) before visualization with a Zeiss Gemini 450 II scanning electron microscope (Carl Zeiss, Ontario, Canada).

#### Bile and acid tolerance assays

The bacteria were cultivated anaerobically in MRS broth for 48h and centrifuged for 10 min at 3500 x *g*. The supernatants were removed, and the bacterial pellets were resuspended and aliquoted in MRS containing 0.3% (w/v) bovine bile (Sigma, Saint Louis, USA) for the bile tolerance assay, and in synthetic gastric juice (pH 3; described by Wall et al) for the acid tolerance assay (48). The bacterial suspensions were incubated at 37°C and samples were taken at 0, 30 and 90 min for the bile acid tolerance, and at 0, 20, 50 and 90 min for the acid tolerance assay. Samples were serially diluted and plated on MRS agar (Merck, Darmstadt, Germany) plates, which were incubated anaerobically for 48 h at 37°C, after which colonies were counted and survival rate calculated.

#### Mucus binding assay

The mucus binding assay performed here was an adaptation of a previous method (49). Briefly, mucus was scraped off from a porcine small intestine, followed by two centrifugations at 11 000 x *g* and 26 000 x *g* for 10 and 15 minutes, respectively. The crude mucus preparation was diluted to OD_280_ 0.1 in PBS and added to each well of a Nunc Maxisorb plate (Nalgene-Nunc, Thermo Fisher Scientific, Rochester, NY, USA) which was then incubated at 4°C overnight under slow rotation. Wells containing mucus were washed three times with PBS (pH 6.0), 0.05% Tween 20 (PBST) and blocked for 60 minutes with PBS + 1% Tween 20 (pH 6.0). The bacterial suspension was diluted to OD_600_ 0.5 followed by two additional washes with PBST after which the bacteria were added to the wells (suspended in PBST (pH 6.0). A timepoint 0 reference was taken before bacteria were added to the wells. The plates were incubated for 4 h at 37°C with slow agitation where after the wells were washed four times with PBST (pH 6.0). Adherent bacteria were treated with trypsin EDTA (0.25%) for 30 minutes at 37°C to detach them from the wells, followed by serial dilutions and plating on MRS plates. The reference sample was also plated. The plates were incubated anaerobically for 48 h at 37°C before viable count was assessed.

#### Antibiotic resistance evaluation

Microdilutions in broth were performed at the Department of Biomedical Science and Veterinary Public Health, Swedish University of Agricultural Sciences, according to ES ISO 10932:2012. Antibiotics tested were Gentamycin, Kanamycin, Streptomycin, Neomycin, Tetracycline, Erythromycin, Chloramphenicol, Ampicillin, Penicillin, Vancomycin, Quinupristin-Dalfopristin, Virginiamycin, Linezolid, Trimethoprim, Ciprofloxacin, and Rifampicin. The control strain *Bifidobacterium longum* ATCC 15707 was used as internal control as described by the ISO method. Reference values were collected from the EFSA guidance on characterization of microorganisms used as feed additives or as production organisms (50).

#### Analysis of biogenic amines and lactate isomers

The analyses were carried out at the Research Institutes of Sweden (RISE). Using 10% inoculum, BG-L47 was inoculated into MRS and an enriched MRS broth containing histidine, lysine, ornithine, and tyrosine (0.25% w/v) and incubated anaerobically at 37°C for 48 h. The bacterial suspensions were centrifugated at 3000 x *g* for 15 min, after which the supernatants were collected and filtered through a 3 kDa filter. Polymeric substances were removed by precipitation using 1% formic acid in acetonitrile, after which the samples were derivatized with AcqTAG and analyzed by LC/MS. Histamine, tyramine, putrescine and cadaverine were quantified against external standards.

For the D- and L-lactate analysis, the solvents were evaporated from the samples which were then derivatized with DATAN-reagent. The labelled lactates were separated and analyzed by LC/MS. D- and L-lactate were quantified against external standards.

#### Sugar metabolism

The *B. longum* strains were cultivated in MRS broth for 48 h under anaerobic conditions at 37°C. The cultures were thereafter streaked on MRS plates (Merck) and incubated for 48 h under anaerobic conditions hours at 37°C. An ampule of API® (bioMérieux, Marcy-l’Étoile, Frankrike) Suspension Medium was opened and a suspension with a turbidity equivalent to 2 McFarland was prepared by transferring a certain number of drops of the bacterial suspension. An ampule of API® 50 CHL Medium was opened and immediately inoculated by transferring twice that number of drops of suspension to the ampule. API 50CH strips were inoculated with the API® 50 CHL Medium and covered with mineral oil. The strips were either incubated for 48 h at 37°C in an anaerobic chamber (anaerobic conditions) or for 48 h at 37°C (aerobic conditions).

### Analysis of glycoside hydrolase genes and fiber utilization

The presence of glycoside hydrolases (GH) was investigated with dbCAN2 (51) using DIAMOND and HMMER. In cases where HMMER did not identify a GH but DIAMOND did, the DIAMOND annotations were considered dominant. For the phenotypic fiber utilization assay, a loop of *B. longum* culture was inoculated into 10 ml of MRS broth and incubated anaerobically at 37°C for 48 h. Modified MRS (mMRS; Supplementary table 3) medium with various polysaccharides (1% w/v) was prepared, 1% *B. longum* culture was inoculated, and the tubes were incubated anaerobically at 37°C for 48 h. Polysaccharides used were fructan (Raftiline HP, ORAFTI s.a. Oreye, Belgium), arabinoxylan (Chamtor S.A., Bazancourt, France), arabinogalactan (Sigma, Saint Louis, MO, USA), β-glucan (Kerry, Tralee, Kerry, Ireland), starch (Sigma cat. 1.01253.500), lacto-N-tetraose (LNT; Glycom A/S, Hørsholm, Denmark), lacto-N-neotetraose (LNnT; Glycom A/S), 3-fucosyllactose (3’FL; Glycom A/S), 2’-fucosyllactose (2’FL; Glycom A/S), 3′-sialyllactose (3’SL; Glycom A/S), 6′-sialyllactose (6’SL; Glycom A/S). mMRS with glucose was used as a control. Final pH was measured after the incubation.

The cellular localization of GH-family enzymes was predicted by analysis with SignalP 5.0 (https://services.healthtech.dtu.dk/services/SignalP-5.0/) and TMHMM (https://services.healthtech.dtu.dk/services/TMHMM-2.0/). The cell surface localization was denoted according to Båth et al. (52). Anchor type was determined by identifying specific domains in the protein sequence.

### Isolation of extracellular MV

*L. reuteri* was co-incubated and grown for 48 h with 25% of bifidobacterial cells (as described in section “Growth stimulation of *L. reuteri* by co-cultivation with *B. longum*”) as a stimulator. After cultivation the bacterial cells were removed in three steps, all at 4°C: *i*) Centrifugation at 4,000 rpm for 10 min, removing the majority of intact cells; *ii*) Centrifugation at 10,000 x *g* for 10 min, removing remaining cells and cell debris; *iii*) Filtration of the supernatant through a 0.45 μm sterile filter (Millipore, Burlington, MA, USA) to ensure that no cells were left in the supernatant.

The MV-containing cell-free supernatant was concentrated using Vivaspin Ultra filtration unit (Sartorius, Göttingen, Germany) with a 100 kDa MwCO until the desired volume of supernatant was achieved. The supernatants were ultracentrifuged at 118,000 × *g* using an 80XP ultracentrifuge (Beckman coulter, Brea, CA, USA) at 4°C for 3 h, followed by resuspension in PBS and ultracentrifuged for an additional 3 h. The pellet was suspended in cell culture medium Neurobasal A or in PBS depending on the subsequent application. The MV preparations were aliquoted and stored at –20°C for later analyses.

The vesicles were sized and quantified using Nanoparticle tracking analysis (NTA). The samples were first diluted to an appropriate volume in PBS and then tracked using NanoSight NS300 (NanoSight^TM^ technology, Malvern, United Kingdom). The tracking was performed with 488 nm blue laser accompanied by a sCMOS camera. The NTA software (version 3.2) was used to record and capture the particles in Brownian motion and calculated the hydrodynamic diameter through the Stokes-Einstein equation, which gives the number of particles per ml.

MV derived from *L. reuteri* DSM 17938 co-cultered with bifidobacterial cells were designated “MV_47_” for BG-L47, and “MV_536_” for BB536 (Table 1). MV derived from *L. reuteri* DSM 17938 grown in SIM with fructose as electron acceptor, without any bifidobacteria, was designated “MV_u_”.

**Table 1:** MV sample names derived from L. reuteri DSM 17938 when B. longum strain was used as stimulant.

| Name | *B. longum* strain |
| --- | --- |
| MV<sub>47</sub> | BG-L47 |
| MV<sub>536</sub> | BB536 |
| MV<sub>u</sub> | None |

### Protein concentration and 5’ nucleotidase activity assays

The protein concentrations of MV preparations were quantified using Qubit protein kit (Invitrogen Carlsbad, CA, USA) according to the manufacturer’s instructions.

The 5’NT activity of the MV preparations was measured using a 5’ Nucleotidase Assay kit (CrystalChem, Elk Grove Village, IL, USA) with slight modifications. Briefly, 10 µL of the samples were added to a 96-well plate containing 180 µL of reagent CC1. After 5 minutes of incubation at 37°C, 90 µL of reagent CC2 was added and the quinone dye was measured at 550 nm after 3, 5, 9 and 13 min. The obtained absorbance differences were used for calculations of 5’NT activities according to the manufacturer’s instructions.

### Evaluation of TRPV1 antagonism

The TRPV1 antagonism assay was performed at Cellectricon, Gothenburg, Sweden. Primary rat dorsal root ganglion (DRG) cultures were extracted from six-week-old Sprague Dawley rats and grown in 384-well plates with the addition of nerve growth factor to mimic peripheral sensitization for two days prior to the experiment. The cells were stained with the FLIPR calcium indicator Calcium 5 Assay Kit (Molecular Devices Corp., Sunnyvale, CA, USA) to image the Ca^2+^-transients evoked by capsaicin addition. One hour prior to the capsaicin/TRPV1 experiments, MV samples were administered at six concentrations (1:3 dilutions in Neurobasal A (Hyclone, GE Healthcare, Chicago, IL, USA) containing Glutamax (Hyclone, GE Healthcare) and supplement B27 (Hyclone, GE Healthcare)). After one hour of incubation with the MV or the reference compound AMG517 (Tocris Bioscience Bristol, UK) transient receptor potential cation channel subfamily V member 1 (TRPV1) evaluations were performed. The 384-well plates were incubated in the Cellaxess Elektra discovery platform (Cellectricon, Gothenburg, Sweden), and the response was continuously measured by capturing changes in the calcium probe fluorescence intensity. The experiments were performed on two separate 384-well plates, using separate compound dilutions. The highest final concentration was 10% of the membrane vesicle stock, corresponding to approximately 10^9^ particles/ml. All concentrations were tested in triplicates and in two biological replicates.

### Secretion of IL-6 and IL-1ß from peripheral blood mononuclear cells

250 000 PBMC were seeded in a 96-well plate at a concentration of 1x10^6^ cells/ml and stimulated with 5x10^8^ MV/ml, corresponding to a multiplicity of MV (MOM) 500, for 48 h at 37°C in 5% CO_2_. The culture supernatants were collected by centrifugation and stored at - 20°C. Secreted levels of IL-1ß and IL-6 were quantified using sandwich ELISA according to the manufacturer’s instruction (MabTech AB, Nacka, Sweden). Absorbance was measured at a wavelength of 405 nm using a microplate reader (Molecular Devices Corp) and results were analyzed using SoftMax Pro 5.2 rev C (Molecular Devices Corp.).

### Whole genome sequencing and genome risk assessment of BG-L47

DNA extraction and preparation of BG-L47 and CF15-4C DNA for whole-genome sequencing was performed as described by Sun et al 2020 (53). Sequencing was performed at the Swedish Veterinary Agency, Uppsala, Sweden, and was accomplished by parallel long read sequencing with Oxford Nanopore (Oxford Nanopore technologies, Oxford, UK) and short read sequencing with Illumina (Illumina Inc., San Diego, CA, USA). DNA for Oxford Nanopore was prepared using the Nanopore Ligation Sequencing kit (SQK-LSK109) with the Oxford Native barcoding genomic DNA kit (EXP-NBD104). The DNA was sequenced using the Oxford Nanopore MinION with a SpotOn flow cell (FLO-MIN106) for 72 h. Base calling was performed using Guppy with the High-Accuracy algorithm. DNA for Illumina sequencing was prepared using the Nextera XT DNA library prep kit (Illumina FC-131-1096) with the Nextera XT v2 Index Kit B (Illumina FC-131-2002) and then subsequently sequenced on an Illumina MiSeq instrument using a MiSeq Reagent kit v3 600 cycle (Illumina MS-102-3033) with a 301 bp read length. The assembly was performed by quality filtering of nanopore reads using Filtlong v0.2.1 (https://github.com/rrwick/Filtlong) and all reads <1 kb were removed as well as 5% of the reads with the lowest quality scores. Illumina reads were quality trimmed using fastp v0.20.1 with the default settings (https://github.com/OpenGene/fastp). Trycycler v.0.5.3 (https://github.com/rrwick/Trycycler) was used for assembly and 12 random read subsamples were generated using trycycler subsample at 75x depth. Each reads subsample was assembled with different assemblers whereas four subsamples were assembled using Flye v2.8.3-b1695 (https://github.com/fenderglass/Flye), four subsamples were assembled using miniasm v0.3-r179 (https://github.com/lh3/miniasm) and minipolish v0.1.2 (https://github.com/rrwick/Minipolish) and four subsamples were assembled using raven v1.4.0 (https://github.com/lbcb-sci/raven). Clusters were subsequently generated using "trycycler cluster". Clusters of good quality were selected manually for downstream analyses whereas other clusters were discarded. Said quality assured clusters were reconciled using "trycycler reconcile". Multiple sequence alignment for the contigs in each cluster was performed using "trycycler msa", raw reads were partitioned between clusters with “trycycler partition”. Using the reconciled clusters, the contig MSA and the partitioned reads, trycycler consensus was used to generate a consensus sequence which were polished using the long read data using Medaka v1.4.4 (https://github.com/nanoporetech/medaka). Finally, the sequences were further polished with the short read data using Polypolish v0.5.0 (https://github.com/rrwick/Polypolish) and POLCA which is part of the MaSuRCA toolkit v4.0.5 (https://github.com/alekseyzimin/masurca). The genome of BB536 (ATCC BAA-999) was readily available at (https://genomes.atcc.org/genomes/).

The genomic risk assessment consisted of mapping of: Clusters of orthologous groups of protein (COG), antibiotic resistance genes, genomic islands (GI) and virulence factors. The mapping of COGs was performed using the eggnog-mapper tool v. 2 (http://eggnog-mapper.embl.de) (54). Settings were left unchanged with exception of SMART annotation. The antibiotic resistance gene search was performed using the CARD database (https://card.mcmaster.ca) and resistance gene identifier (RGI) v. 6.0.1 using the strict setting. In addition, potential acquired resistance genes were searches for using ResFinder v. 4.1 (https://cge.food.dtu.dk/services/ResFinder/). The mapping of GIs and potential GI-borne antibiotic resistance and virulence factors were performed using IslandViewer 4 (https://www.pathogenomics.sfu.ca/islandviewer). The average nucleotide identity (ANI) analysis was mapped using JSpeciesWS ANI calculator (https://jspecies.ribohost.com/jspeciesws/#home) against the reference strains *B. longum* subsp. *longum* ATCC 15707 (KCTC 3128), *Bifidobacterium longum* subsp. *suis* LMG 21814 and *Bifidobacterium longum* subsp. *infantis* NCTC11817.

### Clinical safety and tolerability study

A randomized, double-blind, placebo-controlled clinical study with a parallel-group design was conducted in healthy subjects. The aim of the study was to evaluate safety and tolerability of *B. longum* BG-L47.

The study was conducted by Clinical Trial Consultants AB in Uppsala, Sweden between December 2020 and February 2021. The study was approved by the independent Swedish Ethical Review Authority (Dnr 2020-05404) and registered at ClinicalTrials.gov (NCT04692506) prior the start of the study. The study was conducted in accordance with ethical principles originating in the Declaration of Helsinki (55), and was compliant with International Conference of Harmonization (ICH)/Good Clinical Practice (GCP). Applicable parts of the European Union Clinical Trials Directive and applicable local regulatory requirements were followed. All participants gave verbal and written informed consent before being included in the study. Inclusion and exclusion criteria can be found in Supplementary Tables 4 and 5.

The study was comprised of one screening visit and four study visits: at baseline (day 1), day 7, day 14 and day 28. 36 healthy subjects (24 female) with a mean age of 35.9±13.2 years and a mean BMI of 24.3±3.3 kg/m^2^, were included in the study. Subjects were randomly assigned at a ratio of 1:1:1, to receive treatment low dose (1x10^8^ CFU), high dose (1x10^10^ CFU) or placebo (n=12/group) for a total of 28 days. The study products were sealed one-dose sachets containing lyophilized powder with the strain BG-L47 mixed with maltodextrin. The placebo contained only maltodextrin. Study products sachets were identical, and the contents were similar in appearance. The study product was tested and shown to be stable during storage for the duration of the trial.

The subjects were instructed to mix the content of the sachet with a glass of lactose-free milk before oral consumption in the morning. To be included in the per protocol set, >80% of the study product should have been taken according to instructions without any major deviations that were judged to compromise the analysis of the data. Fecal samples were collected at baseline and after the treatment period at day 28. Vital signs evaluation in the form of systolic and diastolic blood pressure test after 10 minutes of rest was conducted. Physical examination included general appearance including skin, auscultations of lungs and heart, and by abdomen palpation of liver and spleen. Blood samples were collected through venous puncture and *in vitro* assessments of clinical safety were evaluated. This included measurement of chemical parameters such as alanine aminotransferase, alkaline phosphatase, aspartate aminotransferase, bilirubin (total and conjugated), creatinine and glucose (non-fasting). Hematological parameters including hematocrit, hemoglobin (Hb), platelet count and white blood cell count with differential count including leukocytes, lymphocytes, monocytes, neutrophils, eosinophils, and basophils were also evaluated. Follicle stimulating hormone (FSH), serum and urine pregnancy tests were also conducted. To evaluate GI symptoms we used the gastrointestinal symptom rating scale (GSRS), which consists of 15 items that cluster into 5 symptom clades: reflux, abdominal pain, indigestion, diarrhea and constipation (56).

### DNA extraction, PCR and 16S rRNA gene sequencing of fecal samples from the clinical study

Subjects were provided with a sampling instruction manual and instructed to collect stool samples in a sterile stool tube and bring it to the baseline visit (day 1) and final visit (day 28). Briefly, samples of approximately 120 mg were put in Lysis Matrix E tubes (MP Biomedicals) and extracted twice in stool lysis buffer ASL (Qiagen, Hilden Germany). Bead beating at a speed of 5 m/s for 60 seconds (x2) in a FastPrep-24 (MP Biomedicals, Santa Ana, CA, USA) after which each sample was incubated at 90°C and centrifuged for 5 min at full speed at 4°C. Extractions were pooled and subjected to precipitation in isopropanol (volume 1:1). The precipitated DNA was resuspended in TE buffer (VWR, Radnor, PE, USA) and purified using QIAamp DNA Mini Kit (Qiagen). The purified DNA was tested for purity with a nanophotometer (Implen NP80, Munich, Germany), for integrity with a Tapestation 4150 (Agilent, Santa Clara, CA, USA) and the final concentration was determined by fluorometric analysis using the Quant-iT dsDNA HS Assay Kit (Invitrogen) on a Fluoroscan plate reader (Thermo Scientific, Rochester, NY, USA), according to the manufacturers’ instructions.

PCR and 16S rRNA gene amplicon sequencing were performed at NovoGene (Cambridge, UK). PCR to amplify the V4 region was performed using primers 515F (GTGYCAGCMGCCGCGGTAA) and 806R (GGACTACNVGGGTWTCTAAT) (57) ; 20 ng of template DNA was used, and the PCR was limited to 25 cycles (98°C for 1 min, 25 cycles at 98°C (10 s), 50°C (30 s) and 72°C (30 s) and a final 5 min extension at 72°C). The amplicons were sequenced on an Illumina NovaSeq 6000 instrument (paired end reads 250 bp long, 100 K tags per sample). Q30 values varied between 98.30 – 98.79 and the observed OTU’s varied between 611 and 1415 among the samples.

### Strain-specific primer design and qPCR

A short read-based whole genome sequence was acquired for BG-L47 that was draft annotated using RAST (https://rast.nmpdr.org). The genome was blasted against three *B. longum* genomes (*B. longum* DJO10A, *B. longum* NCC2705 and *B. longum subsp. infantis* ATCC 15697), sorted on genes unique for BG-L47 and all sequences longer than 500 bp were translated and analyzed with NCBI BLAST. Unique regions were evaluated, and primer sequences were identified by using Primer3 and Primer BLAST software. Ten 20-bp primers targeting unique regions in a InlB B-repeat-containing protein (accession: “R8G07_03735”) were designed. The primers and qPCR were optimized by running a gradient qPCR with annealing temperatures between 58 and 62°C, determining that 58°C gave the best result.

Primers L47F (ATGGCGATTTCTCCTACCCC) (forward) and L47R (GTACTGGACCATGCGAACCT) (reverse) were able to detect BG-L47 but not BB536 and were chosen for quantification of BG-L47 in fecal samples from the clinical study. The qPCR protocol was 95°C for 3 min followed by 39 cycles of 95°C for 10 s, 58°C for 30 s and 72° C for 10 s.

The log DNA concentration was calculated from Cq-values according to the following calculation, given by the standard curve:

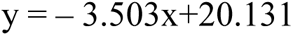

Copy number was calculated using the equation:

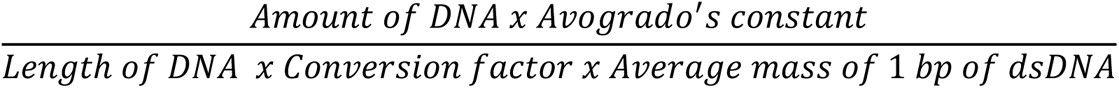

The copy number was then converted to number of bacteria per gram feces with the assumption that the number of copies were equal to number of bacteria. R^2^ value was 0,977 and efficiency was 93%.

### Analysis of microbiota by 16S rRNA gene amplicon sequencing

The DADA2 pipeline (version 1.20, https://benjjneb.github.io/dada2/) was used to clean and filter the raw sequence data and FLASH (58) was used to merge the paired reads. Chimeras were removed and sequences with clean tags were used for subsequent analysis. Amplicon Sequence Variants (ASVs) were generated after being processed by DADA2 to represent de-duplicated unique feature sequences from each sample. After applying QIIME2’s classifier algorithm, each ASV was annotated by a pre-trained scikit-learn Naïve Bayes classifier (58). Based on the annotation of the ASVs, abundances for different bacteria were assigned on kingdom, phyla, class, order, family, genus, and species levels.

The analysis of microbiota composition was done using phyloseq and ANCOMBC packages integrated within the Bioconductor (version 3.14, https://bioconductor.org/) project and vegan (version 2.6, https://github.com/vegandevs/vegan) package. The abundance for all the ASVs were treated by rarefaction method to adjust library differences prior to relative abundance analysis.

The analysis was performed under the R environment (https://www.r-project.org/).

### Statistical analyses

Prism GraphPad version 9.0 (GraphPad Software, Boston, MA, USA) was used in all statistical data interpretations unless otherwise stated. For the growth stimulation, bile tolerance assay, mucus binding assay and TRPV1 antagonism data, ANOVA with Tukey’s *post hoc* statistical analysis was performed. Values below detection limit were set to the detection limit. For the acid tolerance, a mixed effects model with Tukey’s *post hoc* statistical analysis was performed. Paired t-tests were performed for the 5’NT activity analysis. Kruskal Wallis test with Dunn’s multiple comparisons test was performed for the MV protein concentration data. Wilcoxon matched pairs rank tests was performed for the cytokine secretion assay. Statistical analysis of the 16S rRNA gene amplicon sequencing data was performed in RStudio (version 1.4.1717). Alpha diversity was assessed using 4 correction methods: Observed, Shannon, Simpson and InvSimpson. Student t-test was used to compare differences among samples for alpha diversity. Bray-Curtis dissimilarity method was used to assess beta diversity and PCoA was used to reduce dimensionality. Permutational Multivariate Analysis of Variance was used on Bray-Curtis distances for group-wise (HA, HB, LA, LB, PA and PB) and timewise (day 1, 7, 14 and 28) comparisons. A permutation test and the Tukey’s honestly significant difference test (Tukey’s HSD) were then performed on the multivariate homogeneity of group dispersions. Different expression analysis was performed using ANCOMBC and significantly changed ASV feature sequences were then obtained at different taxonomic levels.

For qPCR, the data were baseline corrected and analyzed by ANOVA with Tukey’s *post hoc* test using SAS software version 9.4 (SAS Institute Inc, Cary, NC, USA).

### Data availability

Data not presented in their raw data format are publicly available on Open Science Framework (https://osf.io/jrvq9/?view_only=87d37dd164a749b18b55c40dc429a932). The complete genome of BG-L47 as well as the 16S rRNA genes for BG-L47 and BG-L48 have been uploaded to GenBank with the accession numbers CP137763, MZ411576 and MZ411575, respectively. The plasmid sequence of the mother strain of BG-L47, denoted CF15-4C, has been deposited with the accession number OR817922.

## Results

### Isolation and identification of *Bifidobacterium longum* strains

Seven *Bifidobacterium* isolates from infant feces collected in a study in Tampere, Finland (45), were screened by assessing their antibiotic resistance profiles and cultivability, and two *Bifidobacterium longum* were selected for further characterization (Supplementary table 1). Due to heterogenous colony morphologies when cultivated on MRS agar plates, the two isolates were subjected to repeated cultivation and re-isolation procedures that resulted in strains with stable colony morphology and growth. The strains were named BG-L47 and BG-L48 and deposited to the German Collection of Microorganisms and Cell Cultures (DSMZ) where they were assigned DSM identity numbers DSM 32947 and DSM 32948, respectively. The strains were identified as *Bifidobacterium longum* subsp*. longum* by 16S rRNA gene sequencing (GenBank accession numbers MZ411576 and MZ411575) and by subspecies-specific PCR assays.

### *B. longum* stimulates growth of *L. reuteri* DSM 17938 in simulated intestinal medium

We previously observed that *L. reuteri* DSM 17938 cannot grow in a simplified simulated intestinal medium (SIM; supplementary table 2) without the addition of electron acceptors such as fructose or 1,2-propanediol. The initial aim of the study was to investigate whether specific bifidobacteria could support the growth of *L. reuteri* in SIM without the addition of an electron acceptor. The ability of the two new *B. longum* strains BG-L47 and BG-L48 as well as the reference strain BB536 to support growth of *L. reuteri* DSM 17938 was assessed. Co-cultivation with either of the *B. longum* strains stimulated the growth of *L. reuteri* DSM 17938 (Figure 1) more than the fructose control. The average starting OD_600_ was 0.14 for 4%, 0.26 for 10% and 0.66 for 25% *B. longum* inoculation (Supplementary table 6), and the average final OD_600_ values varied between 0.96 (4% BG-L48) and 2.24 (25% BG-L47). No significant differences were seen between different strains and concentrations (Figure 1). In addition, the co-cultured bacteria were analyzed on HHD diagnostic agar plates, BSM agar and MRS + mupirocin (50 μg/ml) to quantify bifidobacteria and it was revealed that none of the *B. longum* strains had grown in the SIM medium. In fact, all three strains of *B. longum* were reduced more than 100-fold during the incubation period (Supplementary table 7). Thus, the increased OD was a result of *L. reuteri* growth. High performance liquid chromatography (HPLC) analysis showed that supernatants from both *B. longum-*stimulated and fructose-control cultures contained acetate (Supplementary Table 8), indicating that *B. longum* provides *L. reuteri* with an electron acceptor. Metabolites produced by the bifidobacteria were analyzed by nuclear magnetic resonance (NMR) but none of the known electron acceptors of *L. reuteri*, including fructose, glycerol, pyruvate, citrate, nitrate, oxygen and 1,2-propanediol could be detected.

**Figure 1:**
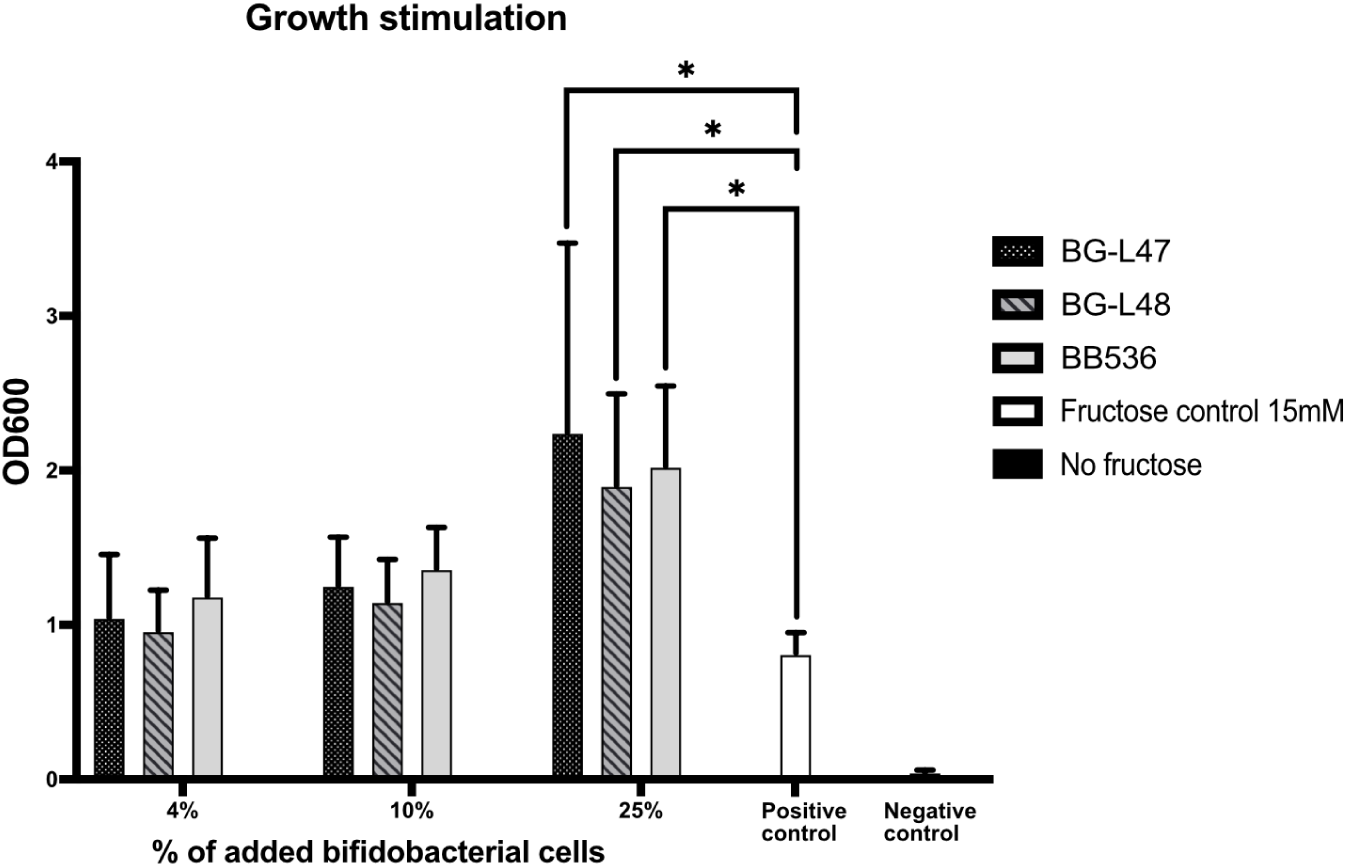
The ability of B. longum strains to stimulate growth of L. reuteri DSM 17938 by co-cultivation in SIM was tested. 4, 10 and 25% of bifidobacterial cell suspension was added and the L. reuteri DSM 17938 cultures were grown for 48 h followed by OD_600_ measurement. Significance level used was *p < 0.05.

### Characterization of the *B. longum* strains

Scanning electron microscopy showed that both BG-L47 and BG-L48 grown in MRS and freeze-dried displayed bulgy morphological appearances. The images reveal smaller particles, which may be vesicular structures appearing on the surface of bacteria as well as in the surrounding space (Figure 2A-B). BG-L47 showed a less bulgy appearance compared to BG-L48. Images of BG-L47 also show larger amounts of web-like structures.

**Figure 2:**
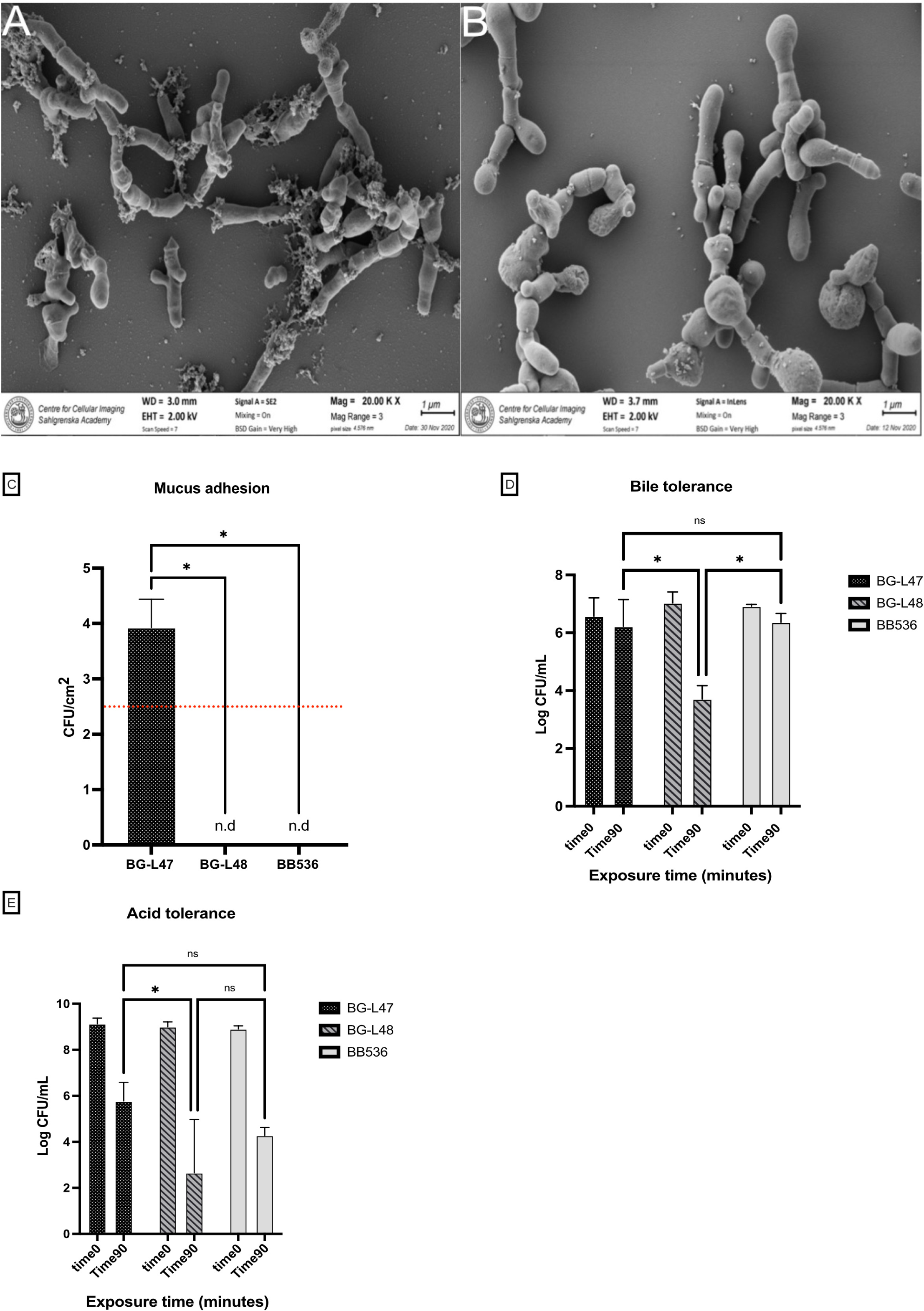
Physiological characterization of the B. longum strains. Scanning electron microscope images showing the morphology of BG-L47 **(A)** and BG-L48 **(B)**. The samples were sputtered with gold before imaging. Adhesion of the B. longum strains to mucus **(C)**. After allowing the strains to adhere to mucus-coated microtiter wells, the numbers of bacteria was determined by trypsin release and plating. Detection limit marked with a dotted red line. Bile tolerance of the B. longum strains **(D)** was determined by exposing the bacteria to 0.3% bovine bile for 90 minutes, after which CFU was measured. Low pH survival of the B. longum strains **(E)** was assessed by exposing the bacteria to synthetic gastric juice at pH 3.0. CFU was measured after 0, 20, 50 and 90 minutes of exposure. The significance level for all statistical analyzes was *p < 0.05.

Physiological properties of relevance to probiotic bacteria were characterized and compared with the reference strain *B. longum* BB536. The mucus binding of the strains was evaluated by quantifying bacteria adhered to a mucus-coated surface. BG-L47 adhered significantly better (*p* < 0.01) than the other two strains, whose adhesion was below the detection limit (Figure 2C). The bile tolerance assay showed that after 90 min of exposure to 0.3% bovine bile, BG-L47 and BB536 had survived more than 2.5 log CFU/ml better (*p* < 0.001) than BG-L48 (Figure 2D). There were no significant differences between the strains at earlier timepoints. In the low-pH tolerance assay performed in synthetic gastric juice at pH 3, BG-L47 decreased by 3 logs after 90 minutes which was 3 logs better than BG-L48 (*p* < 0.01). No significant difference between BG-L48 and BB536, nor between BG-L47 and BB536, was observed (Figure 2E).

The minimum inhibitory concentration (MIC) of a panel of relevant antibiotics was evaluated for BG-L47 and BG-L48. Both strains had MIC values below the cut-off values defined by the European Food Safety Authority (EFSA) concerning susceptibility to antimicrobials of human and veterinary importance (50)(Supplementary table 9).

Sugar fermentation profiles of BG-L47, BG-L48 and BB536 were determined under strictly anaerobic conditions. The results revealed a clear difference between the three strains, as BG-L47 was able to metabolize a broad repertoire of sugars (20 different sugars) and BG-L48 and BB536 were less versatile in their sugar metabolizing abilities (11 different sugars). Tests were performed in five biological replicates and in case of deviations between the results, the fermentation capacity was denoted as variable.

### Identification of glycoside hydrolase genes and analysis of the fiber degradation capacity of BG-L47 and BB536

The results presented so far resulted in the selection of one *B. longum* strain, BG-L47, for further characterization. The strain was further compared with the control strain BB536. A genomic comparison of the presence of glycoside hydrolase genes in available *B. longum* subsp. *longum* genomes revealed that BG-L47 and BB536 are well classified within the traditional clade of *B. longum* subsp. *longum*. BG-L47 encoded a total of 41 glycoside hydrolases (GH) and BB536 encoded 42. BG-L47 did not carry any unique GH compared to BB536 and the three strains DJO10A, ATCC 15707 and NCC 2705 previously described by Vatanen et al. 2022 (59) (Figure 3A). The analysis of glycoside hydrolase genes showed extensive similarities between BB536 and BG-L47, but growth on fiber types associated with the identified GHs differed between the strains. BG-L47 was able to grow on most fibers for which there were one or more corresponding GH gene (except fructan and ß-glucan), whereas BB536 largely didn’t grow on fibers associated with the GH genes in its genome (Figure 3B). A potential explanation could be differences in gene expression between the two strains.

**Figure 3:**
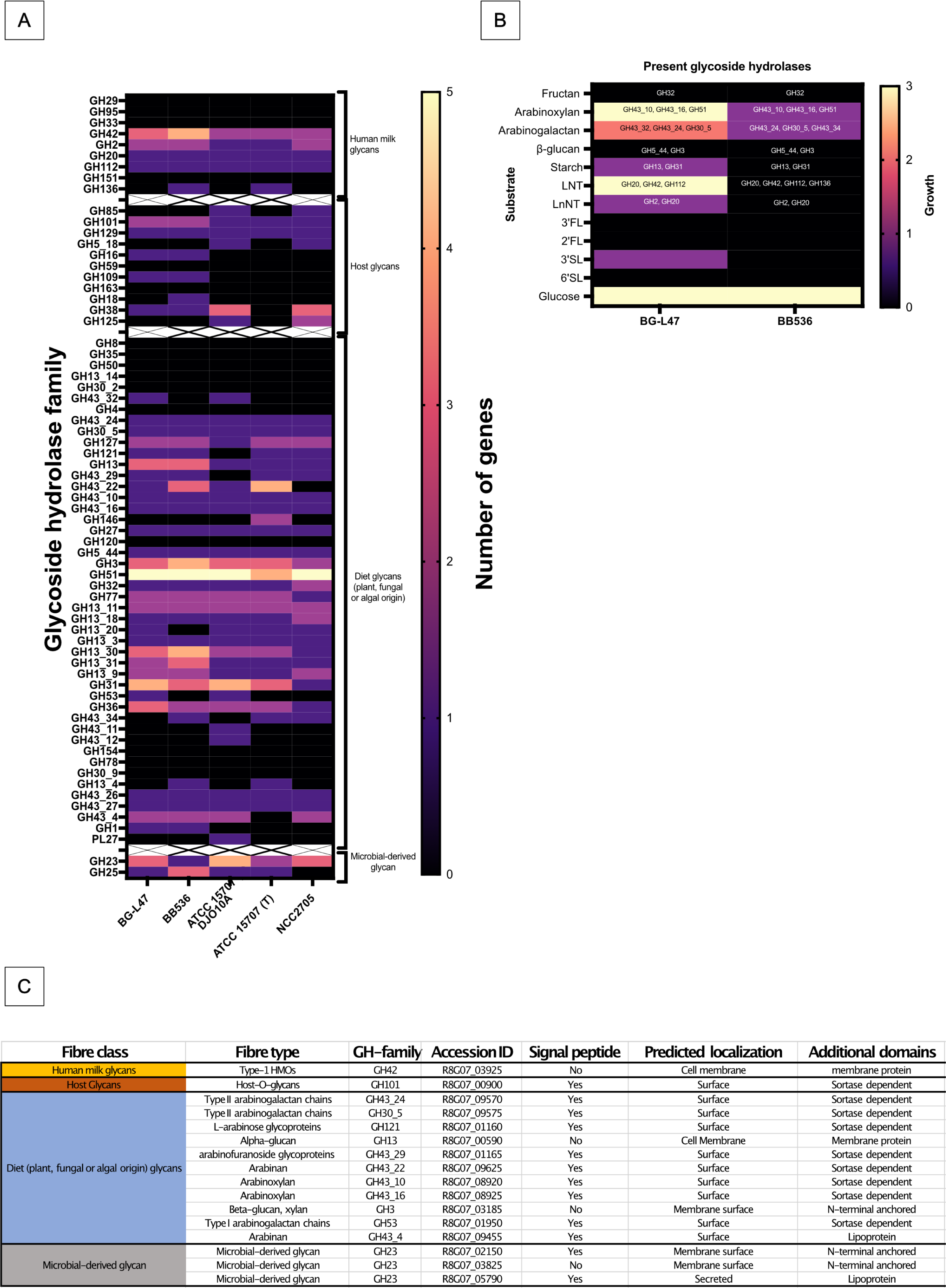
**(A)** Glycoside hydrolases (GH) in B. longum BG-L47 and 4 control strains. The GHs are sorted into four categories according to the origin of the substrate: Human milk glycans, host glycans, dietary glycans and microbially derived glycans. The number of genes within a particular GH family is illustrated by a color scheme representing 0-5. **(B)** Fiber utilization in BG-L47 and BB536 presented together with GHs predicted to be involved in utilization of that specific fiber. Growth was determined by measuring the pH drop when cultured in mMRS with the fibers as carbon source. The explanation of the scale is 0= pH drop 0-0.1, 1= pH drop 0.2-0.5, 2= pH drop 0.5-1, 3= pH drop 1-1.5. **(C)** Predicted extracellular localization of BG-L47 glycoside hydrolases. The protein sequences were analyzed with SignalP, TMHMM, BLAST and subjected to manual annotation; The method for classification of cell surface localization described by Båth et al., 2005 was used.

In BG-L47, 16 of the enzymes were predicted to be secreted or membrane localized (Figure 3C). A majority of the proteins were predicted to be sortase-dependently anchored to the cell wall, but lipoproteins, N-terminal membrane-anchored, and classical membrane proteins were also identified.

### Stimulation of *L. reuteri* DSM 17938 membrane vesicle bioactivity

We have previously shown that extracellular membrane vesicles (MV) can play an important role for the *in vitro* bioactivity of *L. reuteri* DSM 17938 (41,42). In the present study, we have shown that the *B. longum* strains support the growth of DSM 17938, and therefore we investigated whether *B. longum* also stimulated the activity of MV from this strain.

Membrane vesicles were isolated from DSM 17938 grown together with BG-L47 (MV_47_), BB536 (MV_536_) and a control culture with fructose as electron acceptor (MV_u_). The concentrations of the MV preparations were determined to 10^10^ particles/ml by NanoSight analysis. MV_47_ contained ∼5 times as much protein as MV_u_ and ∼2 times as much protein as MV_536_ (Figure 4A). We have previously observed that altered production of MV can be linked to altered activity of 5’ nucleotidase (5’NT) (42), and we therefore tested whether co-cultivation with the *B. longum* strains also affected the 5’NT activity of MV produced by DSM 17938. This analysis showed that MV_47_ had a higher 5’NT activity than both MV_536_ and MV_u_ (Figure 4B).

**Figure 4:**
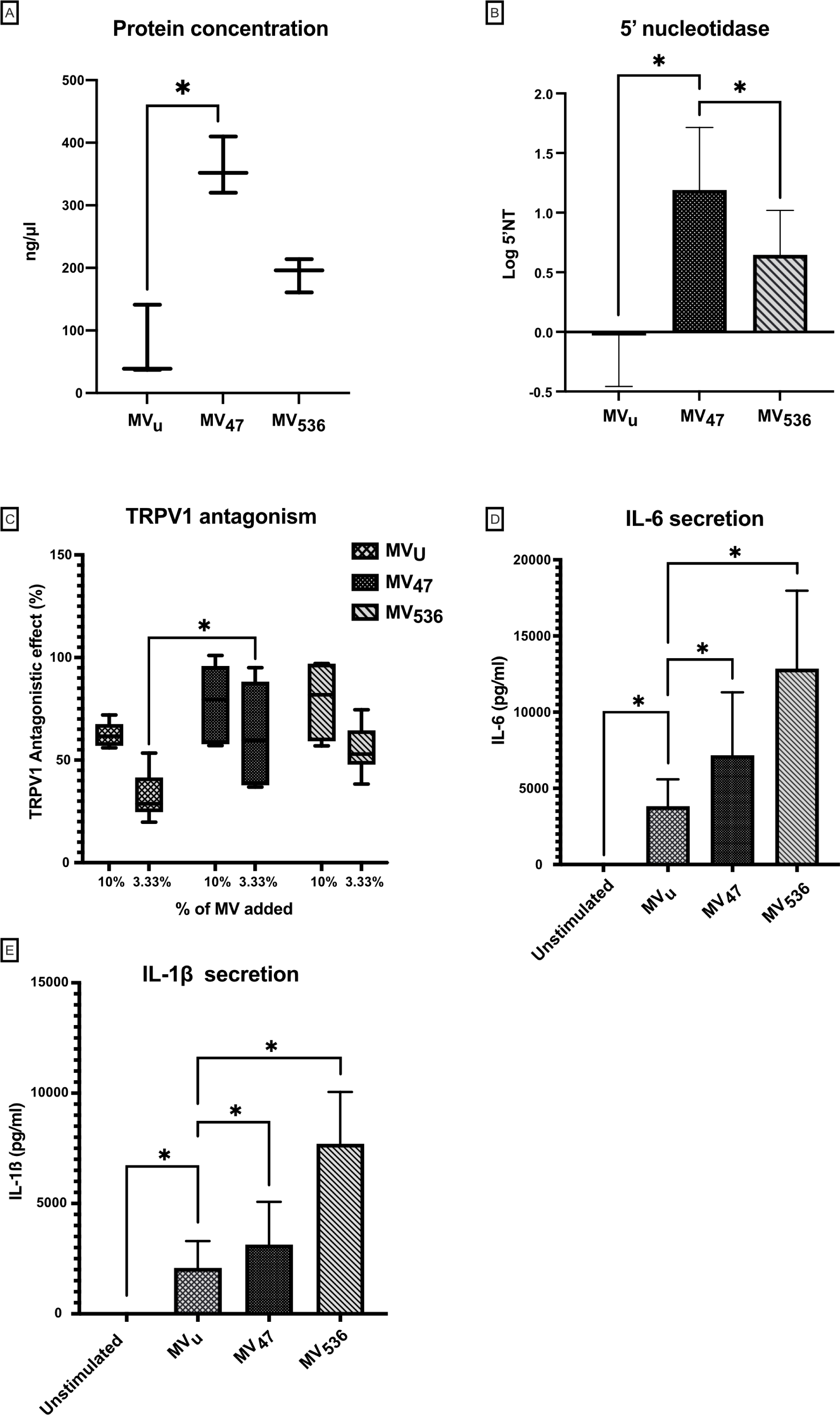
Bioactivities of MV isolated from L. reuteri DSM 17938 co-cultured with B. longum BG-L47 (MV_47_), BB536 (MV_536_), or grown in SIM with fructose as electron acceptor (MV_u_) **(A)** Protein concentrations of MV preparations. Data are presented as boxplots with min to max (n=3). **(B)** 5’NT activity of MV preparations. Values are presented as means with SD (n=5). **(C)** TRPV1 antagonizing effects of MV preparations in a rat dorsal root ganglion (DRG) model. Vesicles were incubated with DRGs, and calcium flux were monitored after addition of the agonist capsaicin, n=6. Boxplots with min to max and average value representation. **(D)** and **(E)** MV stimulation of IL-6 and IL-1ß secretion in PBMCs from healthy donors measured by ELISA (n=6). The significance level in all graphs is *p < 0.05.

It is known that *L. reuteri* and its MV can antagonize the TRPV1 receptor on dorsal root ganglia and stimulate IL-6 and IL-1ß in naïve PBMCs (41,42). In this study, all vesicle preparations including MV_u_ showed antagonistic effects on TRPV1 upon capsaicin stimulation but MV_47_ had a higher antagonistic effect than MV_u_ (Figure 4C). In addition, all MV preparations stimulated the production of IL-6 and IL-1ß, but MV_47_ and MV_536_ induced a stronger response than the control vesicles MV_u_ (Figure 4D-E).

### Genome Characteristics and Safety Assessment of BG-L47

The genome sequencing of *B. longum* BG-L47 resulted in a closed chromosome and no plasmid was identified. The genome consists of 2378960 base pairs, has a GC content of 60.1% and a total of 1941 coding sequences. The average nucleotide identity (ANI) analysis showed that BG-L47 is phylogenetically closest to subsp. *longum*. The ANI scores for the different subspecies were 98.1% for *B. longum* subsp. *longum* ATCC 15707, 96.3% for *B. longum* subsp. *suis* LMG 21814 and 94.4% for *B. longum* subsp. *infantis* NCTC11817 (Supplementary table 10). The genetic comparison between BG-L47 and CF15-4C revealed that a cryptic plasmid of 3624 base pairs had been lost during the selection procedure (plasmid accession number OR817922).

The genome safety assessment showed that BG-L47 have a COG profile comparable to BB536 (Supplementary table 11). There was no production of any biogenic amines or of D-lactate (Supplementary table 12). The total number of assigned COGs in BG-L47 was 1565, which corresponds to 80.6% of the whole genome. BB536 had a total of 1556 COGs assigned, corresponding to 76.9% of the genome. Interestingly, compared to BB536, BG-L47 encodes an additional 11 proteins belonging to the COG group G (carbohydrate transport and metabolism) in accordance with its broader sugar utilization capacity. None of the strains carry any antibiotic resistance gene. BG-L47 has 45 genomic islands (GI) and BB536 has 48, and there were no gene encoding resistance or virulence factors on these (Supplementary table 13). In conclusion, there were no safety concerns in the genome of *B. longum* subsp. *longum* BG-L47.

### Clinical Safety and Tolerability Study of BG-L47

Finally, we evaluated BG-L47 in a clinical safety and tolerability study. There were no clinically significant differences in any either group in the primary endpoints vital signs, physical examination and frequency, intensity, or seriousness of adverse events (AE) (Table 2). There was one severe AE in the low-dose group, but after analysis by the clinical investigator it was deemed unlikely to be related to the treatment. Blood samples showed no significant differences in hematocrit, hemoglobin, platelets, and leukocytes at day 28 compared to day 1. The total mean of gastrointestinal symptom rating score (GSRS) showed no clear differences between either treatment group at any of the timepoints (day 1, 7, 14 and 28). Likewise, there were no clear differences in the mean total gastrointestinal symptom rating score over time within the treatment groups. Of 36 participants, 32 had 100% compliance, meaning that the study product was consumed every day as directed. Four subjects, three in the placebo group and one in the high-dose group, missed one dose each. In conclusion, BG-L47 at a dose of up to 10^10^ CFU/day was safe and well tolerated during the study.

**Table 2:** API of B. longum BG-L47, BG-L48 and BB536 under strictly anaerobic conditions. Growth was assessed by color change which was estimated manually on a scale of 0-5 * Indicates that the results varied between the five biological replicates.

|  | BG-L47 | BG-L48 | BB536 |
| --- | --- | --- | --- |
| **Total** | 20 | 11 | 11 |
| L-arabinose | 5 | 5 | 5 |
| D-ribose | 1* | 5 | 0 |
| D-xylose | 5 | 0 | 5 |
| D-Galactose | 5 | 5 | 5 |
| D-Glucose | 5 | 5 | 5 |
| D-Fructose | 5 | 2 | 5 |
| D-Mannitol | 5 | 0 | 0 |
| D-Sorbitol | 5 | 0 | 0 |
| Methyl-α-D-glucopyranoside | 1* | 0 | 0 |
| N-Acetylglucosamine | 3* | 0 | 0 |
| Arbutin | 3* | 0 | 0 |
| D-Maltose | 5 | 5 | 5 |
| D-Lactose (bovine origin) | 5 | 5 | 5 |
| D-Melibiose | 5 | 5 | 5 |
| D-Saccharose | 5 | 5 | 5 |
| D-Melezitose | 5 | 0 | 0 |
| D-Raffinose | 5 | 5 | 5 |
| Starch (Amidon) | 1* | 0 | 0 |
| Glycogen | 1* | 0 | 0 |
| D-Turanose | 3 | 0 | 3 |
| Potassium gluconate | 0 | 3 | 0 |

**Table 3:**
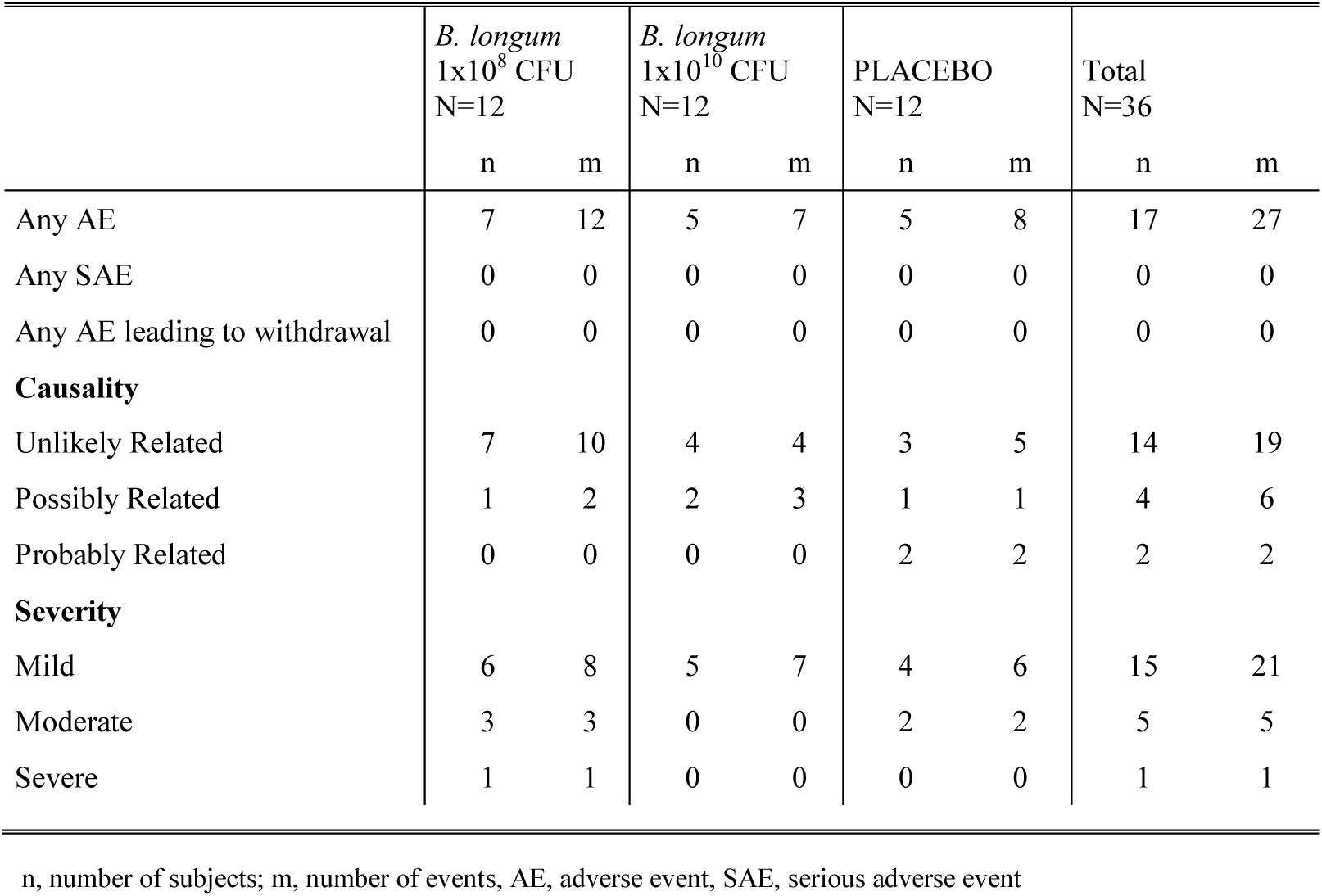
Overview of adverse events.

|  | *B. longum*<br>1x10<sup>8</sup> CFU<br>N=12 |  | *B. longum*<br>1x10<sup>10</sup> CFU<br>N=12 |  | PLACEBO<br>N=12 |  | Total<br>N=36 |  |
| --- | --- | --- | --- | --- | --- | --- | --- | --- |
|  | n | m | n | m | n | m | n | m |
| Any AE | 7 | 12 | 5 | 7 | 5 | 8 | 17 | 27 |
| Any SAE | 0 | 0 | 0 | 0 | 0 | 0 | 0 | 0 |
| Any AE leading to withdrawal | 0 | 0 | 0 | 0 | 0 | 0 | 0 | 0 |
| **Causality** |  |  |  |  |  |  |  |  |
| Unlikely Related | 7 | 10 | 4 | 4 | 3 | 5 | 14 | 19 |
| Possibly Related | 1 | 2 | 2 | 3 | 1 | 1 | 4 | 6 |
| Probably Related | 0 | 0 | 0 | 0 | 2 | 2 | 2 | 2 |
| **Severity** |  |  |  |  |  |  |  |  |
| Mild | 6 | 8 | 5 | 7 | 4 | 6 | 15 | 21 |
| Moderate | 3 | 3 | 0 | 0 | 2 | 2 | 5 | 5 |
| Severe | 1 | 1 | 0 | 0 | 0 | 0 | 1 | 1 |

n, number of subjects; m, number of events, AE, adverse event, SAE, serious adverse event

The effect of the intervention on the microbiota composition was investigated by 16S rRNA gene amplicon sequencing of fecal samples from the clinical study. The analysis showed that there were no significant changes in alpha diversity at the phylum and family levels (Figure 5A, data not shown). At the genus level, alpha diversity scores were significantly reduced in the high-dose group with Simpson, inverted Simpson, and Shannon correction (Supplementary Figure 1). Interestingly, the dispersion of beta diversity at the genus level (supplementary table 14) showed that there was a strong time-dependent effect that was significant for all comparisons between pre- and post-intervention groups. This indicates either an overtime effect or a matrix effect.

**Figure 5:**
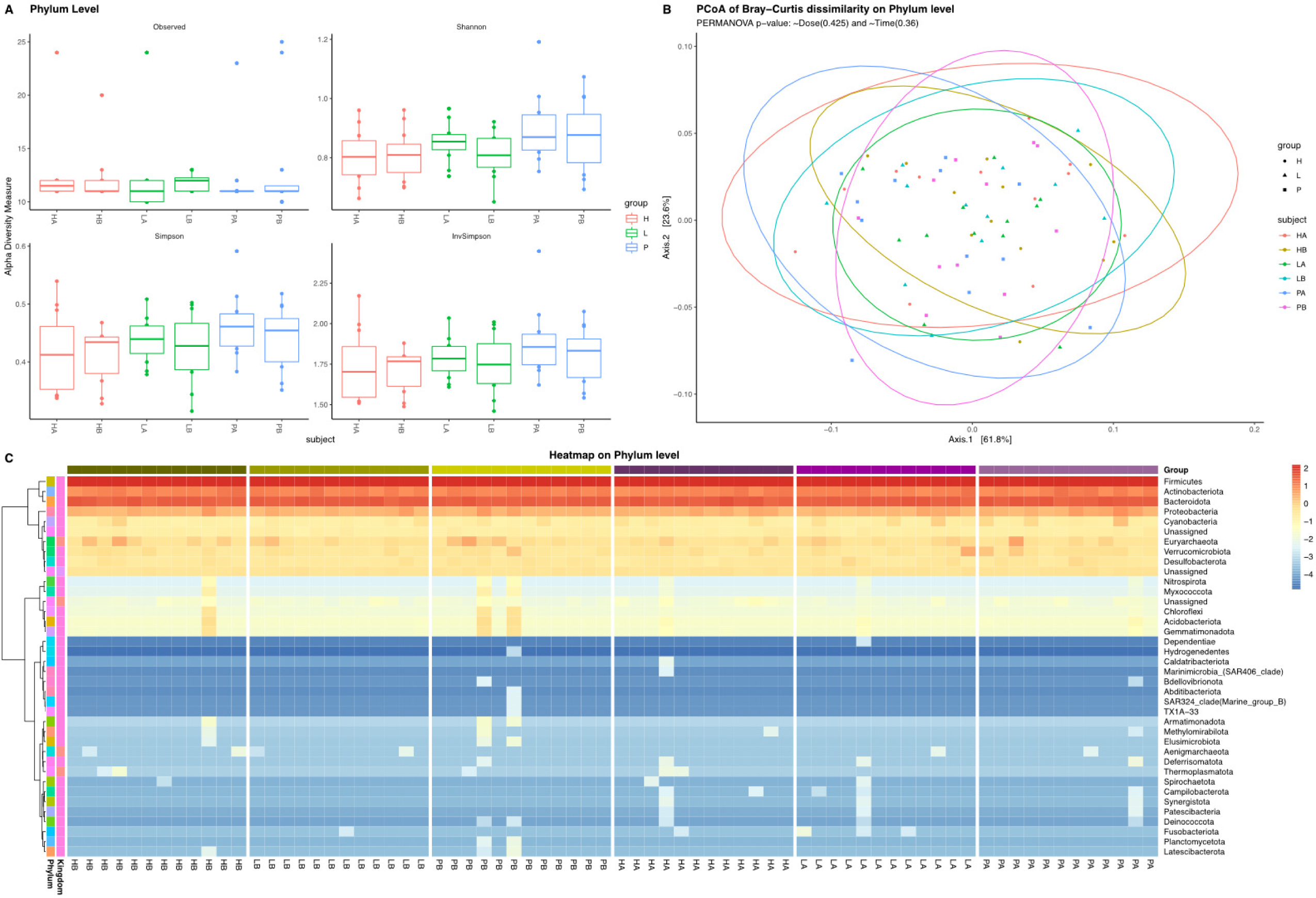
**(A)** Alpha diversity at the phylum level. **(B)** Principal coordinate analysis (PCoA) plot of treatment groups at the phylum level. **(C)** Heatmap showing the relative abundance at phylum level for all participants. Data are shown as log relative abundance adjusted by their individual mean. HA denotes high dose after treatment, HB denotes high dose group before treatment, LA denotes low dose after treatment, LB denotes low dose group before treatment, PA denotes placebo group after treatment, PB denotes placebo group before treatment.

The microbial composition at the phylum level was found to be similar when comparing day 1 to day 28 (Figure 5B). There were no clear differences in relative abundance at the group level, although some participants differed (Figure 5C). A total of 25 different genera changed significantly after treatment with the high-dose product (Supplementary table 15). Further investigation revealed that the genera that decreased in abundance in a majority of participants in the high-dose group were *Solobacterium* (*S. moorei*) and *Gracilibacteraceae* (unknown species). However, the most distinct difference was an increase of *Odoribacter* (*O. splanchnicus*) observed among all participants in the high-dose group (Supplementary Figure 2). None of these genera were significantly changed in the placebo or low-dose groups. There were 19 genera changed in the low dose group and 20 in the placebo group (Supplementary table 15). Finally, no apparent increase in *Bifidobacterium* or *Lactobacillus* was observed when assessing the relative abundance.

The presence of BG-L47 was quantified by qPCR targeting a strain-specific gene encoding a protein containing InlB B repeats and the levels detected corresponded well with the doses administered (Table 4). The median number of BG-L47 (or naturally occurring similar strains) after 28 days of intervention was 7.7 log_10_ CFU g^−1^ for the high-dose group, 5.5 log_10_ CFU g^−1^ for the low-dose group and 2.9 log_10_ CFU g^−1^ for the placebo group. At day 0, all three groups had 2.0 – 2.7 log_10_ CFU g^−1^. Two participants in the placebo group had detectable levels of the target gene at both timepoints (6.2 to 9.0 log_10_ CFU g^−1^). ANOVA with baseline correction and Tukey’s adjustment showed that the high-dose group had significantly more BG-L47 compared to the low-dose and placebo group. The low-dose group had significantly more BG-L47 compared to the placebo group.

**Table 4:** Median fecal levels of B. longum BG-L47 in participants receiving either high or low dose BG-L47 or placebo. Values are median (interquartile range) of samples shown in log_10_ CFU g^−1^. * Denotes significance against both other groups on day 28. ANOVA with Tukey’s adjustment and baseline correction.

|  | Day 0 | Day 28 |
| --- | --- | --- |
| **BG-L47 High dose** | 2.0 (0.0–3.3) | 7.7 (7.1–8.2)<sup>*a</sup> |
| **BG-L47 Low dose** | 2.7 (0.0–3.1) | 5.5 (4.6–6.8)<sup>*b</sup> |
| **Placebo** | 2.3 (0.0–3.4) | 2.9 (2.5–3.9)<sup>*</sup> |

- a. High dose vs low dose: (estimate=2.1335 (95% CI 3.7704; 0.4966), $p \leq 0.01$ ), high dose vs placebo: (estimate=4.9144 (95% CI 6.5709; 3.2579), $p \leq 0.0001$ ) b. Low dose vs placebo: (estimate=2.78 (95% CI 1.12; 4.44), $p \leq 0.001$ ).

## Discussion

*Lactobacillaceae* and *Bifidobacteriaceae* are two abundant bacterial families in the infant gut and are also common in adults. Both bacteria have been associated with multiple health benefits and incidence-preventive effects (29,60). Although there are many commercial probiotic products on the market containing multiple strains that lack support for synergistic interactions, we sought to investigate a potentially synergistic effect of combining two species that may encounter each other in the gut. Bifidobacteria and lactobacilli have been shown to co-localize in the human neonatal gut and may likely interact there (61). By growing them in a substrate with similarities to the content of the intestine (Simulated Intestinal Medium, SIM), we aimed to mimic some of the interactions that occur in the gut with a potential benefit for the probiotic activity. Combinations of lactobacilli and bifidobacteria have been explored by others but bacterial interactions in an intestinal-like environment, resulting in both increased growth and bioactivity, has not, to our knowledge, been previously studied.

We first showed that strains of *B. longum* stimulated the growth and later also the bioactivity of *L. reuteri* DSM 17938 membrane vesicles. Previously, Adamberg et al. 2014 showed that specific strains of *Bifidobacterium* and *Lactobacillus* could grow synergistically after passage through a gastrointestinal tract simulator (62). We found that the addition of *B. longum* provides an electron acceptor required by *L. reuteri* to grow in SIM. This type of synbiotic relationship has been previously described and the authors showed that 1,2-propanediol, a potential electron acceptor for *L. reuteri*, was produced and secreted by *Bifidobacterium breve* and then increased the growth of *L. reuteri* (9). The electron acceptor produced by the bifidobacteria in this study was not one of the known electron acceptors of *L. reuteri* (fructose, glycerol, pyruvate, citrate, nitrate, oxygen and 1,2-propanediol) and the identity of the molecule remains unknown.

Survival in the gastrointestinal tract is a desirable probiotic trait and therefore tolerance to gastric pH and bile was evaluated. BG-L47 showed the strongest tolerance profile of the tested strains. Tolerance to these stressors has previously been evaluated for *B. longum* BB536, a strain capable of deconjugating bile salts (63), and our results show that BG-L47 is as tolerant as BB536. Analysis of the genome sequence of BG-L47 revealed that the strain carries a bile salt hydrolase gene (cholylglycine hydrolase; EC 3.5.1.24; “R8G07_04930”). It has previously been shown that BB536 is sensitive to acidic pH (63), an observation that was valid for all three *B. longum* strains tested in our study. The sensitivity to acidic pH in *B. longum* (and many other bifidobacteria) in contrast to *B. animalis* has been suggested to originate in the inability of the susceptible species to upregulate the activity of H^+^-ATPase (64), making them unable to pump out enough protons in this stressful environment. However, here we show that BG-L47 tolerates an acidic environment better than BG-L48, making it a potential better probiotic strain candidate. Furthermore, adhesion to the mucosa is a requirement for transient colonization of the intestine (65) and the mucus layer represents one of the most prominent sites for microbe-host interactions (66). Interestingly, we show that BG-L47 was the only strain for which mucus binding could be detected. In another type of adhesion assay, BB536 has been shown to bind HT-29 epithelial cells (63). However, intestinal epithelial cells are covered with mucus and in a healthy intestine bacteria usually do not come in direct contact with the epithelium and therefore the ability to adhere to mucus may be a prerequisite for effective interactions with the host.

A versatile carbohydrate metabolism is an important characteristic for bacterial colonization of the intestine (23,67), and interestingly, there was a clear difference in the number of sugars that BG-L47 could metabolize (20 in total) compared to BB536 and BG-L48 (11 in total). Degradation of polysaccharides is a hallmark of bifidobacteria, and BG-L47 and BB536 encode a total of 41 and 42 glycoside hydrolases, respectively, which is comparable to a number of other *B. longum* subsp. *longum* strains (Figure 3A). Vatanen and colleges (59) refer to transitional *B. longum* subsp. *suis/suillum* as strains that can utilize both substrates present in breast milk and in solid foods of plant origin. Also, BG-L47 can utilize substrates of breast milk and plant origin. Interestingly, BB536 encodes multiple HMO-associated GHs but was unable to grow on any of the HMOs, while BG-L47 encodes fewer GH families of the HMO utilization cluster but grew well on LNT. It has previously been described that BB536 is unable to ferment HMOs (68), although LNT was not evaluated. Both BG-L47 and BB536 are genetically equipped with an ABC transporter operon predicted to be involved in LNT, lacto-N-biose (LNB) and galacto-N-biose degradation. In BG-L47 the transporter is encoded by R8G07_08360, R8G07_08365, and R8G07_08370, and by an ortholog of the operon in BB536 (HCBKAFLH_01753, HCBKAFLH_01754, HCBKAFLH_01755).

However, the inability of BB536 to grow on LNT indicates that the ortholog is not expressed or unable to transport LNT, a phenomenon that has been observed in other *B. longum* strains with orthologs of the *Blon_2177-2175* operon (69). Yamada and colleagues further showed that *B. longum* subsp. *longum* JCM1217, which has an ortholog with 99% sequence identity to the BB536 *Blon_2177* ortholog (HCBKAFLH_01755), is unable to transport LNT. Instead, the orthologs were shown to transport only LNB released from LNT degradation by the extracellular GH136 lacto-N-biosidase LnbX, an enzyme also found in BB536 (69). Overall, the reason for the absence of growth of BB536 on LNT is not known but may be due to expression of LNT metabolism associated genes.

Another interesting finding was that several GH enzymes had motifs typical of sortase-dependent cell-wall anchored proteins. Those motifs have been described as indicators of important host interaction proteins, often involved in adhesion (70,71). Sortase-dependent cell-wall anchored enzymes involved in fiber degradation on the surface of *B. longum* BG-L47 potentially indicates an ecological role in facilitating cross-feeding in the intestine by degrading fibers into less complex sugars that can be used by other intestinal inhabitants. Similarly, extracellular degradation of xylans has been described in *Bifidobacterium pseudocatenulatum,* emphasizing the role bifidobacteria as primary fiber degraders in the human gut (72). All three *B. longum* strains were relatively tolerant to oxygen and could to grow under semi-aerobic conditions (substrate covered by oil) but exhibited variable sugar metabolism profiles when grown under these conditions (supplementary table 16). The apical mucosa closest to the lumen maintains approximately 0.1-1% oxygen diffusing into the lumen (73) but the intestine of a newborn is known to be more oxygenated than that of adults (74). Thus, a *B. longum* strain with a favorable tolerance profile and a versatile carbohydrate metabolism could be ecologically favored in the infant gut.

*L. reuteri* DSM 17938 expresses and secretes 5’-nucleotidase (5’NT), an enzyme that converts AMP to the potent signal molecule adenosine (75). We have previously used the activity of 5’NT to evaluate biological activity in MV (42). Interestingly, the lactobacilligenic effect of BG-L47 was stronger than that of BB536 because the 5’NT activity in *L. reuteri* DSM 17938 MV stimulated by BG-L47 (MV_47_) was significantly higher than when stimulated with BB536 (MV_536_). Similarly, we observed that the antagonistic effect on the pain receptor TRPV1 increased significantly in response to MV_47_. The increased 5’NT activity in MV_47_ could potentially partially explain the increased antagonistic effect on TRPV1 as inhibitory interactions of adenosine on TRPV1 have been suggested previously (76). TRPV1 interactions have many clinical implications (77) including IBS and abdominal pain (78), atopic dermatitis (79), airway inflammation (80) and potentially infantile colic (81). TRPV1 is also involved in shaping the immune system (82) as well as the microbiota (83) and may be a key site of interaction for further exploration. In contrast, MV_47_ induced a slightly lower cytokine secretion from PBMC compared to MV_536_. This is interesting because adenosine causing IL-6 induction has been reported by others (84), suggesting that it is not adenosine alone that causes IL-6 release from PBMCs in this model. Early colonization and increased abundance of *Bifidobacterium* early in life has been correlated with an increased immune responsiveness and the ability to produce higher levels of both IL-6 and IL-1β (27).

Qualified presumption of safety (QPS)-listed *B. longum* BG-L47 was well tolerated by healthy adults with no clinically significant differences in adverse events, vital signs, physical examinations, or biochemical markers and no differences in GSRS scores between groups given a high dose BG-L47, low dose BG-L47 or placebo. Analysis of fecal samples showed that the number of BG-L47 found by strain-specific qPCR corresponded well with the high- and low-dose treatment groups as well as the placebo treatment. The 16S rRNA gene sequencing revealed no differences at higher taxonomic levels. However, at the genus level there were differences in alpha diversity in the participants who received the high dose of BG-L47. High diversity is often considered beneficial, but it has previously been described that fiber intervention can result in reduced alpha diversity (85). Possibly, interventions with potent fiber degraders or just fibers may produce similar effects on microbial diversity. As a final note, larger studies would be required to verify the findings in altered alpha diversity. There were changes in the composition in all groups at the genus level, and in the high dose group we observed a reduction of *Solobacterium moorei*, which has been described as a pathogen that can be a cause of halitosis and invasive infections such as bacteremia (86,87). *Gracilibacteraceae*, a commensal in the oral cavity (88), was also reduced in the high-dose group. Another interesting observation in the high-dose group at the genus level was the increased relative abundance of *Odoribacter splanchnicus*, a bacterial species that produces acetic acid, propionic acid, succinic acid, and low levels of butyric, isovaleric and isobutyric acids. It is a common inhabitant of a healthy microbiota and interestingly, *O. splanchnicus* and its MV have shown anti-inflammatory effects on the gut epithelium (89). At the genus level, there were also differences in beta diversity between all groups before and after treatment, suggesting that either the intervention vector, i.e., glass of milk or maltodextrin, or/and the time of the study, influenced the microbiota composition. The study was conducted between December and January, a time when many people change their nutritional habits.

We have focused on MV secreted from *L. reuteri* DSM 17938, but upon co-incubation of the B. longum strains with *L. reuteri* there is a possibility that some of the MV were released from the bifidobacteria. To address this, we performed an additional experiment in which *B. longum* was incubated in SIM for 48 h. From these cultures we extracted MV and measured their 5’NT activity. No pellet could be seen and no 5’NT activity was detected, indicating that the bioactivity of the cocultures was indeed derived from *L. reuteri*. Evaluation of interactions within bacterial consortia is very complex but increased understanding of MV may be important for future studies of these types of interactions (35).

In summary, the search of a partner strain that stimulates *L. reuteri* resulted in the selection of BG-L47. The strain was initially selected for its processability, superiority in tolerances and mucus binding, and ability to utilize a broad range of carbon sources. In addition, BG-L47 has the ability to boost the activity of *L. reuteri* DSM 17938 and its derived MV. The strain was considered safe, but future clinical studies should investigate the probiotic effect in specific groups of subjects. In addition, more research is needed to elucidate the mechanisms by which *B. longum* affects *L. reuteri* or acts as a stand-alone probiotic strain.

## Acknowledgments

We would like to thank Jonas Faijerson Säljö and Pernilla Garberg for their support during this work, Hans Jonsson (Department of Molecular Sciences, Swedish University of Agricultural Sciences) for valuable input and discussions, Helena Bysell for her active involvement in the early stages of the project, Cellectricon for providing the TRPV1 model, Anna Pielach at the Center for cellular imaging (University of Gothenburg) for aiding with SEM imaging. We further thank Roger Karlsson (Nanoxis Consulting AB), Anders Karlsson (Nanoxis Consulting AB) and Beatriz Piñeiro Iglesias (Department of Infectious diseases, Sahlgrenska academy, University of Gothenburg) for fruitful discussions and preparations before the SEM imaging, Erik Rein-Hedin (CTC Clinical Trial Consultants AB) for supervising and aiding with the clinical safety study, Simon Isaksson (SLU) for help with the HPLC analysis and Aldina Pivodic (APNC Sweden AB) for help with statistical interpretations.

